# Conformational Reorganisation of Apolipoprotein E Triggered by Phospholipid Assembly

**DOI:** 10.1101/2020.08.18.255299

**Authors:** Dube Dheeraj Prakashchand, Jagannath Mondal

**Affiliations:** Tata Institute of Fundamental Research, Center for Interdisciplinary sciences, Hyderabad 500107, India

## Abstract

Apolipoprotein E (ApoE), a major determinant protein for lipid-metabolism, actively participates in lipid transport in central nervous system via high-affinity interaction with lipoprotein receptor LDLR. Prior evidences indicate that the phospholipids first need to assemble around apoE, before the protein can recognise its receptor. However, despite multiple attempts via spectroscopic and biochemical investigations, it is unclear what are the impact of lipid assembly on globular structure of apoE. Here, using a combination of all-atom and coarse-grained molecular dynamics simulations, we demonstrate that, an otherwise compact tertiary fold of monomeric apoE3 spontaneously unwraps in an aqueous phospholipid solution in two distinct stages. Interestingly, these structural reorganizations are triggered by an initial localised binding of lipid molecules to the C-terminal domain of the protein, which induce a rapid separation of C-terminal domain of apoE3 from the rest of its tertiary fold. This is followed by a slow lipid-induced inter-helix separation event within the N-terminal domain of the protein, as seen in an extensively long coarse-grained simulation. Remarkably, the resultant complex takes the shape of an ‘open conformation’ of lipid-stabilised unwrapped protein, which intriguingly coincides with an earlier proposal by small-angle X-ray scattering (SAXS) experiment. The lipid-binding activity and the lipid-induced protein conformation are found to be robust across a monomeric mutant and wild-type sequence of apoE3. The ‘open’ complex derived in coarse-grained simulation retains its structural morphology after reverse-mapping to all-atom representation. Collectively, the investigation puts forward a plausible structure of currently elusive conformationally activated state of apoE3, which is primed for recognition by lipoprotein receptor and can be exploited for eventual lipid transport.

Table of Content

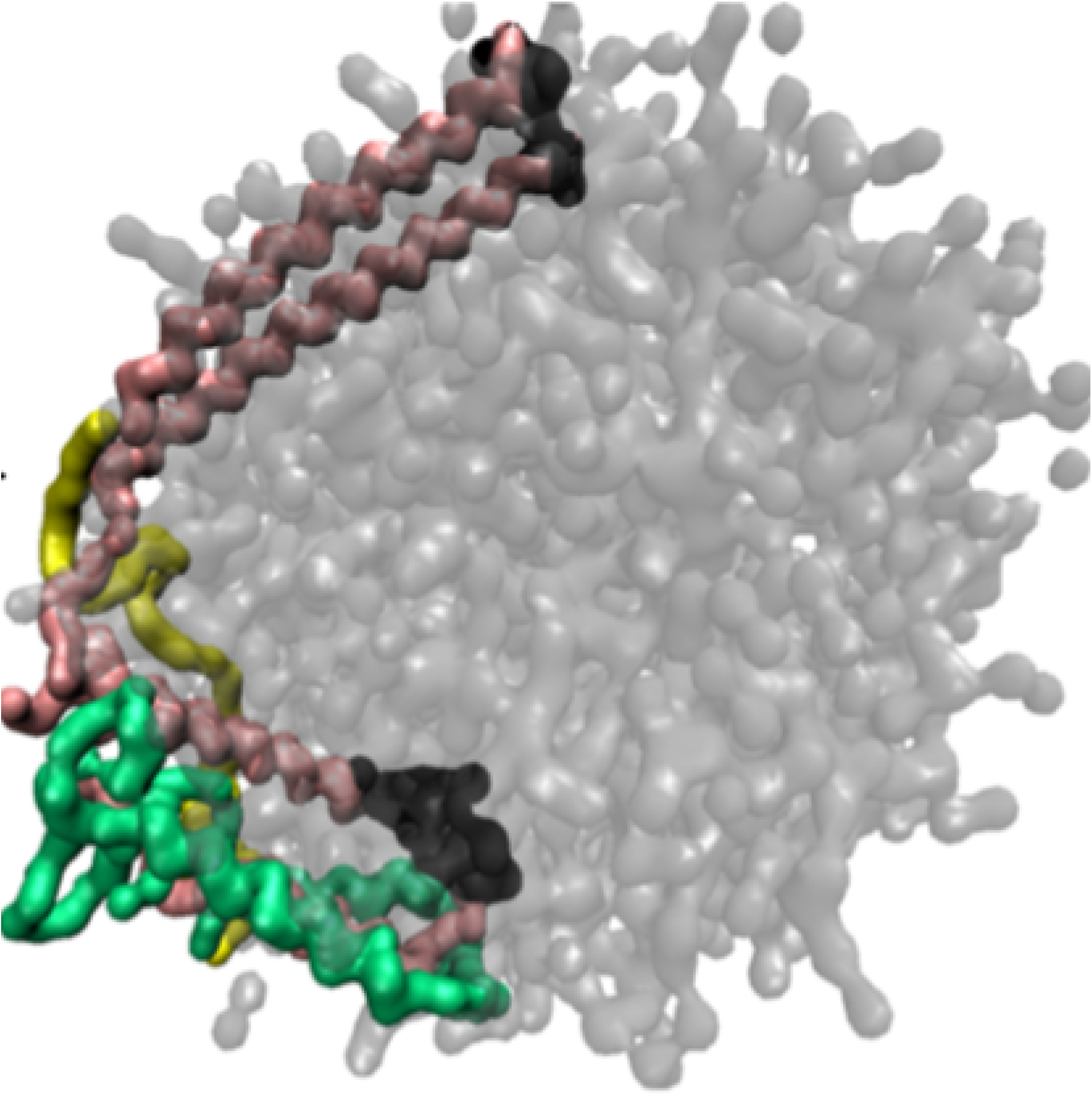

## Introduction

The intense hydrophobic character of triglycerides and cholesterol prevents them from getting dissolved in the aqueous environment of the blood. These hydrophobic entities are transported in blood plasma in the form of large assemblies of lipids (like phospholipids, triglycerides, cholesterol-esters and free-cholesterol) and proteins called lipoproteins.^1,2^ Each of these lipoproteins carry different proteins serving as address tags which decide the destination and hence the function of each lipoprotein complexes. Apolipoprotein E (ApoE), a 299-residue protein consisting of N-terminal domain (NTD) and C-terminal domain (CTD) connected by a hinge-region,^3–5^ constitutes the protein component of lipoproteins not only in plasma^6^ but also in the central nervous systems (CNS)^7–10^ and serves as one of the major determinants in lipid metabolism.^11^ As a part of its lipid-transport activity, low-density-lipoprotein receptor (LDLR) recognises apoE as a ligand. However, in the absence of lipid molecules, apoE is not recognised by LDLR. Prior experimental investigations indicate that as a prerequisite for apoE to bind to the LDLR,^12,13^ apoE needs to be conformationally activated via complexation with lipid.^14–17^ It has also been demonstrated that the complexation of this protein with phospholipids gives rise to a apoE.lipid lipoprotein particle that binds efficiently to the LDLR.^18^ More over, full-length apoE has been found to bind to LDLR on fibroblasts only after complexation with lipid.^19^ All these circumstantial evidences point to the fact that the receptor recognition properties of this protein are manifested only in a lipid-associated state. However, a molecular-level insight into the mechanism associated with apoE-lipid complexation process and plausible morphology of the resulting lipoprotein particle are currently elusive.

Over the years, model lipoproteins composed of apoE and phospholipids have served as a suitable invitro mimic of many of the critical biological activities of naturally occurring apoE-containing lipoproteins.^20^ Phospholipids, including dimyristoylphosphatidylcholine (DMPC) or dipalmitoylphosphatidylcholine (DPPC) have been used to create nanometer scale lipoprotein particle wherein the apoE circumscribes the periphery of a disk-shaped phospholipid bilayer.^20,21^ On the basis of biophysical and spectroscopic studies, various models of apoE-phospholipid complexes have been proposed.^4,21–23^ These models basically suggest possible ways in which the N-terminal helix bundle may change its structure upon lipid association. Small angle X-ray scattering (SAXS) based investigations of morphological features of apoE-DPPC lipoprotein complex have proposed that these complexes are ellipsoidal in shape and the morphology of the phospholipid core is compatible with a twisted-bilayer model.^20^ However, a separate investigation has crystallized apoE-DPPC complex at a 10 angstrom resolution and predicted an incomplete toroidal model.^24^ While the general consensus is that the protein undergoes a lipid binding-induced conformational change, the ultimate conformation adopted by apoE in presence of phospholipids is still a matter of debate. This is mostly due to the lack of clarity about the nature of a molecular-level conformational change that apoE undergoes in the process of lipid binding.

Recently, a series of biophysical and spectroscopic studies have brought out interesting features about the interactions of apoE and phospholipids.^25–29^ As a key observation, Garai and co-workers, using fluorescence-based kinetic studies, have illustrated that while all lipid-free isoforms can exist primarily as monomers, dimers and tetramers at micro molar concentration, lipid binding takes place only with the monomeric form of apoE.^26^ It has also earlier been inferred that monomeric form of apoE prevails on lipoprotein surface.^27^ The report of a high-resolution NMR structure of the full-length monomeric mutant of apoE3 by Chen et al^30^ (see figure 1a) has rejuvenated the call for a molecular-level hypothesis on apoE3-lipid interaction and associated conformational organisation that the apoE3 might possibly undergo upon lipid association. By combining existing hydrogen-deuterium exchange data and via inferring from NMR-derived structure and non-equilibrium computer simulations, Frieden and coworkers^31^ have proposed possible mechanisms of conformational organisation of apoE3 via lipid binding in between N-terminal and C-terminal domain.

**Figure 1:**
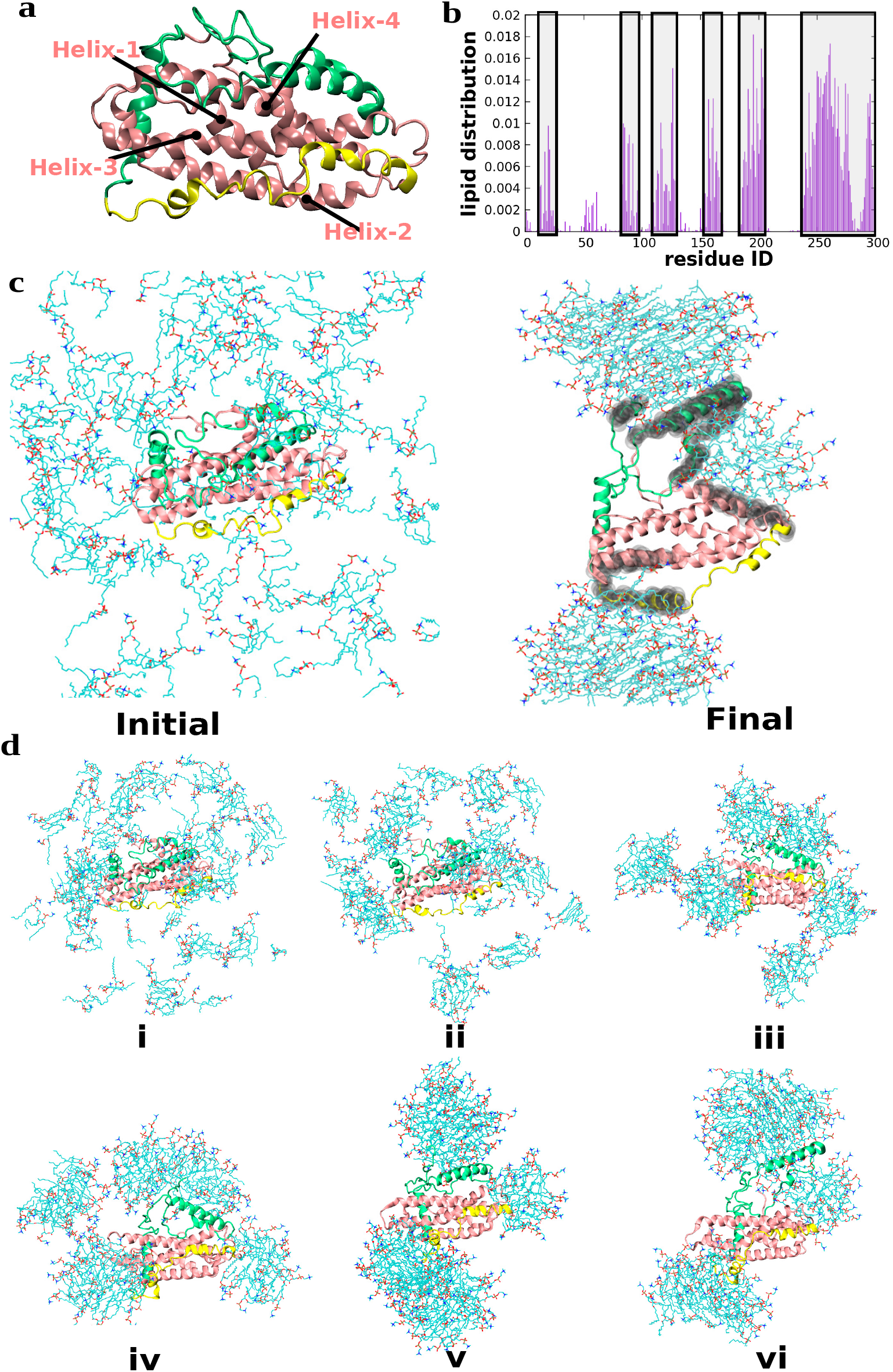
**a.** NMR-derived structure of apoE3 with N-terminal (NT) domain marked as pink; the hinge region in yellow; and the C-terminal (CT) domain shown as green. The four helices of NT domain helical bundle are also pointed out. **b**. Normalised probability of distribution of lipids around the residues of apoE3. **c**. Left: protein in lipid-dispersed state in the beginning of the simulation and Right: protein in the lipid-equilibrated phase of the simulation near the end of 2 *µs*-long simulation. Water not shown for clarity. Color code for 7 DPPC lipid: hydrophobic tail marked in cyan and polar headgroups marked in red color. **d**. All-atom simulation snapshots show the spatial distribution of the lipids around the protein at different time points of the simulation.

However, a vast range of investigations notwithstanding, an atomic-level picture of the dynamical process of lipid-induced conformational organisation associated with apoE.phospholipid complexation process is yet to emerge. The current work closes this gap by capturing the phospholipid binding-induced conformational change of E3 isoform of the protein (hereby referred as apoE3) via extensive molecular dynamics simulations. Specifically, by utilising both all-atom and coarse-grained model we report a high-resolution dynamical picture of apoE3-DPPC complexation process in explicitly modelled water at two distinct stages. First, we show that the C-terminal domain of monomeric apoE3^30^ moves away from N-terminal domain in presence of DPPC lipids within the period of our atomistic MD simulation. The atomistic simulations identify the important regions of protein where the lipids would mostly settle. The trajectories elucidate the key local changes in the helicity, salt-bridges and intrinsically disordered regions of apoE3 as a consequence of lipid binding. In the subsequent step, via adopting a coarse-grained model, we demonstrate the slow *on-the-fly* unwinding of N-terminal domain upon lipid association. The resultant conformation provides a high-resolution picture of the lipoprotein complex. The final equilibrated lipid-stabilised morphology of apoE3 in the lipoprotein takes the shape of a lipid-stabilised ‘open conformation’, previously hypothesized in small-angle X-ray scattering experiments.

## Methods and Materials

The investigation involved molecular dynamics simulation in two different stages. The first stage involved simulation using atomistic models and the final stage involved simulation using coarse-grained model. All MD simulations described in the current work were performed with the Gromacs 2018 simulation package^32^, in most cases benefiting from usage of Graphics processing units (GPU).^33^ To check the robustness of the simulation results across different software platform we have also repeated some of the simulations using 2020 version of Gromacs.

### Atomistic Simulations

In our work, all simulation involved single copy of E3 isoform of apoE (referred as apoE3) in its monomeric state. The NMR-derived structure of monomeric apoE3 (figure 1a) by Chen et al(PDB: 2L7B)^30^ is a mutant form of the wildtype apoE3 where five hydrophobic residues of C-terminal domain had been mutated to polar residues (F257A/W264R/V269A/L279Q/V287E). This mutant form of apoE3 is previously well-known for its ability to remain in monomeric form upto a reasonably high concentration (5-20 mg/mL).^34,35^ More importantly, past experiments have demonstrated that structure and stability of this quintessential monomeric mutant are identical to wild-type apoE3.^34,35^ Finally, both the mutant and wild-type apoE3 are known for displaying similar lipid-binding activities.^35^ Accordingly, due to presence of well-resolved NMR structure, the monomeric mutant formed the basis of current investigation. Nonetheless, we have also performed additional simulations involving wild-type protein for further validation of our results (after back-mutating the NMR-derived structure). DPPC lipid molecules were randomly distributed around apoE3. 135 lipid molecules were required to fill the simulation box. 49862 TIP3P^36^ water molecules are used to solvate the simulation box and the system was rendered charge-neutral by replacing five water molecules by equal number of potassium ions. A periodic box was implemented with the box dimension of the equilibrated system in each direction equal to 11.9 nm. The protein, lipid and ions were modelled using charmm36^37,38^ force field. The system is first energy-minimised using steepest descent minimisation scheme.

The energy minimization step is then followed by an equilibration for 100 ps in NVT ensemble at an equilibrium temperature of 320 K using the Velocity rescale thermostat^39^ with time constant 0.1 ps. The system is then equilibrated for 500 ps in NPT ensemble at equilibrium temperature of 320 K using the same thermostat with a time constant 1 ps and the equilibrium pressure of 1 bar using the Parrinello-Rahman^40^ barostat with time constant 2 ps using isotropic pressure coupling. All hydrogen bonds were constrained using LINCS^41^ algorithm and the water molecules were kept rigid using SETTLE algorithm.^42^ Verlet cutoff scheme was employed in all simulations. The short-range interactions were modelled by Lennard Jones interaction with a cutoff length of 1 nm and with dispersion correction. We have also checked the robustness of the simulation results by repeating additional simulations via turning off dispersion corrections. The long-range electrostatics was handled using Particle mesh Ewald (PME) scheme via a cubic interpolation and Fourier spacing of 0.16 nm and a real space cutoff of 1 nm. Six realisations of system, differed by initial velocities, was then propagated for at least 2 *µ*s using leap-frog algorithm with an integration time step of 2 femtosecond. As a control simulation, a lipid-free simulation of apoE3 in neat water was performed using similar protocol. Towards this end, six trajectories are performed for 1.5 *µs*, with randomised initial velocity, in aqueous medium which contained 31267 water molecules with a box size of 9.867 nm and maintaining a temperature of 320 K.

### Coarse-grained Simulations

The final protein conformation of all-atom simulated trajectories, in which the C-terminal domain has moved apart from the rest of the apoE3, was chosen for coarse-grained mapping and long time simulations. The coarse-grained protein is lipidated with 135 DPPC lipid molecules. MARTINI coarse-grained model^43–45^ was used as the force field of choice. The periodic boundary condition was employed and the empty spaces within the box was solvated by 12204 MARTINI non-polarisable water. The equilibrated box dimension was 11.86 nm. The system was charge-neutralised as well. The Verlet cut-off scheme used was for the simulation. The short-range nonbonding interaction was modelled by Lennard Jones interaction with a short-range cutoff of 1.1 nm. As per Martini protocol, the electrostatics was modelled via reaction field with a short-range coulomb cutoff of 1.1 nm and dielectric constant of 15. The system was subjected to energy minimisation and NPT equilibration. Thermostat used were velocity-rescale^39^ for both equilibration and production run while the barostat used was berendsen for equilibration and parrinello-rahman^40^ for production run. Two realisations of 55*µs* long MD simulation, starting with independent initial configurations, were employed to evolve the system maintaining a temperature of 320 K, with an integration time step of 30 fs.

## Results

The precedent experimental investigations have established that the apoE3 binds with phospholipid in its *monomeric* form.^26^ Accordingly, the current computer simulations focus on elucidating the mechanism of lipid complexation with monomeric apoE3. As would be revealed in our investigations, we demonstrate a stage-wise kinetic molecular picture of overall complexation process :

- In a first rapid step, lipid induces separation of C-terminal domains of apoE3 from the remaining fold.
- In the final step, a slow unwinding of N-terminal domain of apoE3 takes place. In the current article, we employ all-atom and coarse-grained molecular dynamics simulations respectively, to access the relevant time scale associated with these two steps and provide a molecular view of the spontaneous apoE3-lipid complexation process.

### Lipid induced separation of C-terminal domain of ApoE as the first step

The NMR derived structure of monomeric apoE3 by Chen et al^30^ forms the basis of our work for exploring its mechanism of interaction with lipid molecules (see figure 1a). Following previous works,^30,31^ we define residues 1-167 as N-terminal domain, residues 206-299 as C-terminal domain and residues 168-205 as the hinge helix region. We note that the NMR structure was derived by Chen et al^30^ after mutating the wild type apoE3 in five residues (F257A/W264R/V269A/L279Q/V287E) of C-terminal domain, to circumvent the issue of the protein’s self-oligomerization. These mutations helped retain the monomeric state which ultimately allowed for recording efficient and accurate NMR structure. In our present investigation the NMR structure of this monomeric mutant forms the basis of the study, due to availability of the high resolution structure and past experimental demonstration of its similarity with wild-type apoE3 in the context of structure, stability and lipid-binding activity.^34,35^ Accordingly, most of the investigations in this work involve monomeric mutant and its interaction with lipid particles. Nevertheless, as would be discussed later, we have also performed additional simulations with backmutated wild type apoE3 in presence of lipids for a validation of the proposed mechanism of lipid-induced conformational change in monomeric mutant.

### Localized lipid-assembly around apoE3 C-terminal domain

The monomeric apoE3 in its all-atom representation is solvated with water and 135 DPPC lipid molecules are randomly placed inside a cubic box of 11.9 nm. The left panel of figure 1c depicts a snapshot of the initial configuration of the system. Multiple realisations of initial configuration of the system, thus prepared, are individually subjected to multi-microsecond long equilibrium MD simulations at temperature of 320 K. As depicted in supplemental movie S1, we find that lipid molecules diffuse from the bulk media and assemble around the protein during the course of these microsecond-long simulations. Left and right panels of figure 1c illustrate the representative snapshots of equilibrated conformation of apoE3 in presence of DPPC lipids, as obtained at the beginning and at the end of 2 *µ*s long MD simulation respectively. We find that, starting from a dispersed state in bulk water, the lipid molecules eventually get settled mostly near the C-terminal domain of the monomeric apoE3. The spatial density map of the lipid molecules around protein, averaged over the simulation trajectories, also indicates a localised accumulation of lipid molecules near the C-terminal domain (see grey coloured density map around C-terminal domain in figure 1c, right). Figure 1b plots the statistical propensity of lipid accumulation within 0.5 nm cutoff distance of each constituent residue of apoE3, averaged over six simulation trajectories, each at least 2 *µ*s long. Consistent with spatial density profile, we find that large segments of C-terminal domain (residues 243-296) are mostly surrounded by lipids, while there are also isolated locations around N-terminal domains (residues 13-24 , 84-97, 108-130, 186-206) showing non-negligible lipid density. This result is consistent with previous experimental reports of preferential lipid binding propensities by C-terminal domain of apoE3.^25,28,29^ We find that the C terminal domain presents a large exposed hydrophobic surface which acts as the initiator of interactions with lipids. The preferential binding of lipid molecules to the C-terminal domain of apoE3, as captured in these all-atom simulations, also resonates very well with the proposed hypothesis by Chen et al based on their NMR structure of monomeric apoE3.^<u>30</u>^

### Lipid-induced separation of C-terminal domain of apoE

As an intriguing feature of the snapshot represented in the right panel of figure 1c, significant structural reorganisation of apoE3 tertiary fold due to phospholipid accumulations is clearly evident, especially relative to its initial lipid-dispersed state (figure 1c, left). Specifically the snapshot indicates that C-terminal domain of the apoE3 has moved away from the rest of the protein’s tertiary fold in presence of lipids. For comparison purpose, figure 2 depicts representative equilibrated snapshots of apoE3 in neat water with that in the presence of phospholipid. Consistent with our earlier finding,^46^ we find that the tertiary fold of apoE3 remains intact in neat water, in contrast to the unravelling of C-terminal domain of the same protein in lipid-water mixture. For more statistically rigorous comparison, SI figure S1 provides snapshots of the protein in the two cases (neat water versus lipid-water mixture) obtained near the end of six independent trajectories, each at least 1.5 *µs* long. This comparison confirms that the conformational reorganisation of apoE3, as observed in the current simulation trajectories is caused due to the presence of phospholipids and is not merely a result of thermal fluctuation. Figure 1d provides a time evolution of lipid assembly process, with the snapshots i-vi rendering the representative snapshots at different time points. We find that the lipid molecules, initially in randomly solvent-dispersed state, gradually undergo spontaneous self-assembly process and take the shape of micelles. Some of these micelles coalesce and simultaneously approaches the protein surface. The coupled process of micellization and their protein adsorption eventually result in the localised accumulation of lipid-assembly predominantly around C-terminal domain of the protein and certain isolated locations in the N-terminal domains (discussed earlier). The supplemental movie S1 captures the dynamical process of conformational organisation that apoE3 undergoes in presence of lipid during 2 *µs*-long all-atom MD simulations. We find that C-terminal domain of apoE3 gradually moves away from the helix 3/helix4 interface of the N-terminal domain within microsecond-long time scale in majority of our simulation trajectories. On the other hand, the tertiary fold of N-terminal domain remains largely unperturbed within the time scale of our all-atom simulations. This lipid-induced structural change in protein, elucidated in right panel of figure 1c and figure 2, warrants a detailed molecular-level characterisation, as would be elaborated in the upcoming sections.

**Figure 2:**
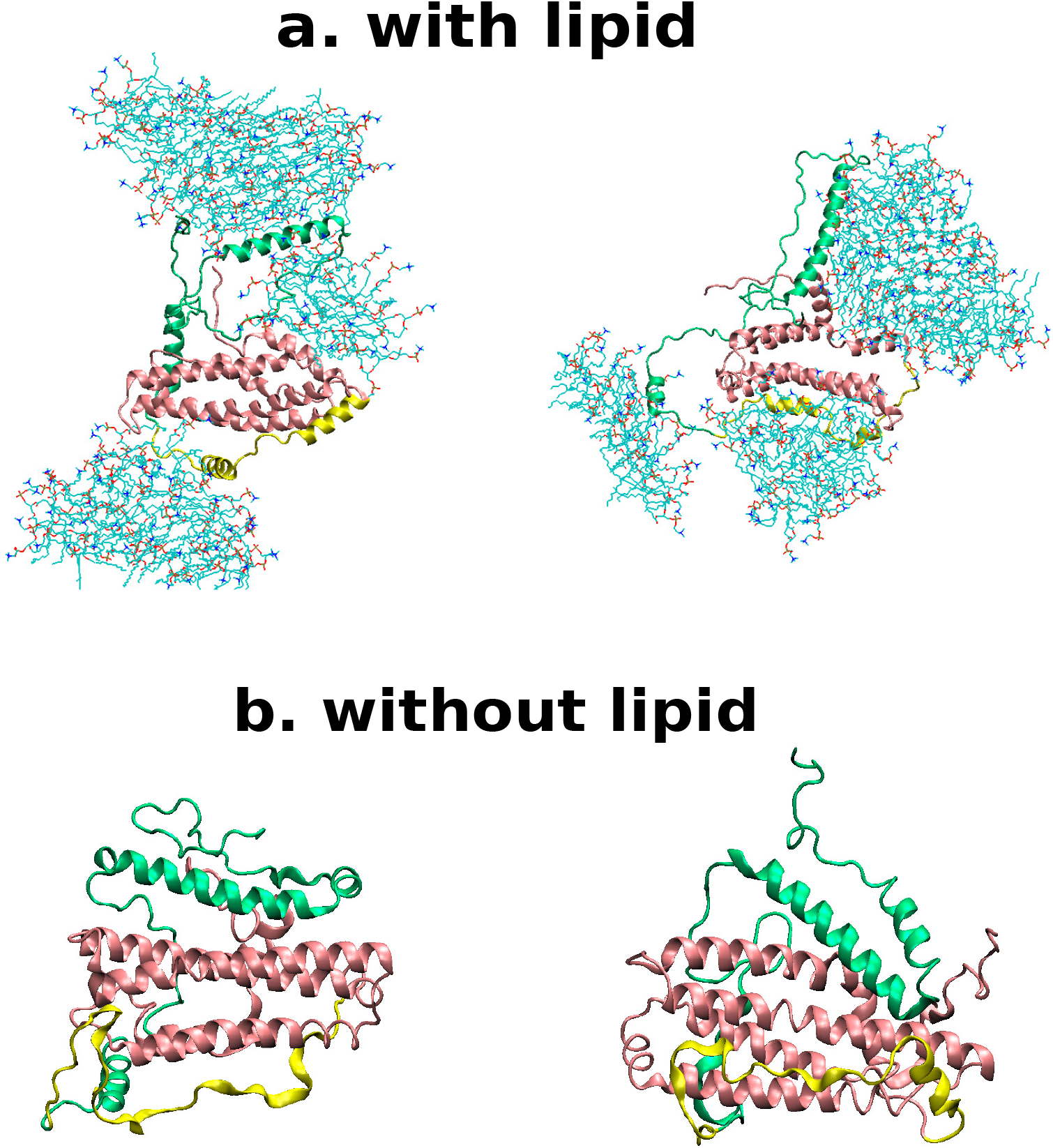
**a.** The representative snapshots obtained in two independent all-atom trajectories with simulation box containing 135 molecules of DPPC along with the water molecules (water not shown). **b**. The structures attained in two independent all-atom trajectories without any lipids molecules. Both types of simulations were run for at least 1.5 *µs*. (refer to the color code described in the figure 1). Refer to figure S1 for comparison across more trajectories

As a result of assembly of lipids around C-terminal domain of the apoE3, we find that the C-terminal domain of the protein gradually starts separating from the rest of the protein. Figure 3a represents the time profile of root-mean-squared deviation (RMSD) of C-terminal domain, relative to the NMR-derived structure of apoE3. In line with the supplemental movie S1, we find that RMSD of the C-terminal domain (figure 3a) steadily increases with time and plateaus to a large value of 2-2.5 nm in majority of the MD simulation trajectories. The simulation-snapshots in figure 3c, extracted from key time-points of RMSD profile of C-terminal domain, illustrate the gradual progression of the unwinding process of the C-terminal domain of apoE3 away from N-terminal domain. On the other hand, the RMSD of the N-terminal domain, relative to NMR structure, is relatively muted and ranges between 0.5-1 nm (figure 3b). Together, this analysis is consistent with the event of preferential localised lipid accumulation around the C-terminal domain (see figure 1c). This observation from our simulation also supports earlier hypothesis^30,31^ that N-terminal domain acts as the folding template of C-terminal domain in the NMR-derived structure of apoE3. We find that the number of C-terminal domain atoms within 0.5 nm radial distances of Helices 3/4 of N terminal domain acts as a sensitive parameter for characterising the extent of opening of the C-terminal domain in presence of lipid. As seen in this time profile of figure S2, the number of neighbouring C-terminal atoms around (within a radius of 0.5 nm) the helix 3/4 of the N-terminal domain gradually reduces with time. As we would see in the later part of the article, this lipid-induced separation of C-terminal domains, away from the N-terminal domain, would provide an opening for eventual unwinding of N-terminal domain. Thus, the contacts lost between the C-terminal and N-terminal domains would be replenished by the contacts between lipids and the N-terminal domain. The overall picture of lipid-induced unwinding of the protein is robust across different version of simulation programs (Gromacs 2020 vs 2018) (figure S3) and holds true irrespective of incorporating the dispersion corrections for nonbonding interactions (figure S4).

**Figure 3:**
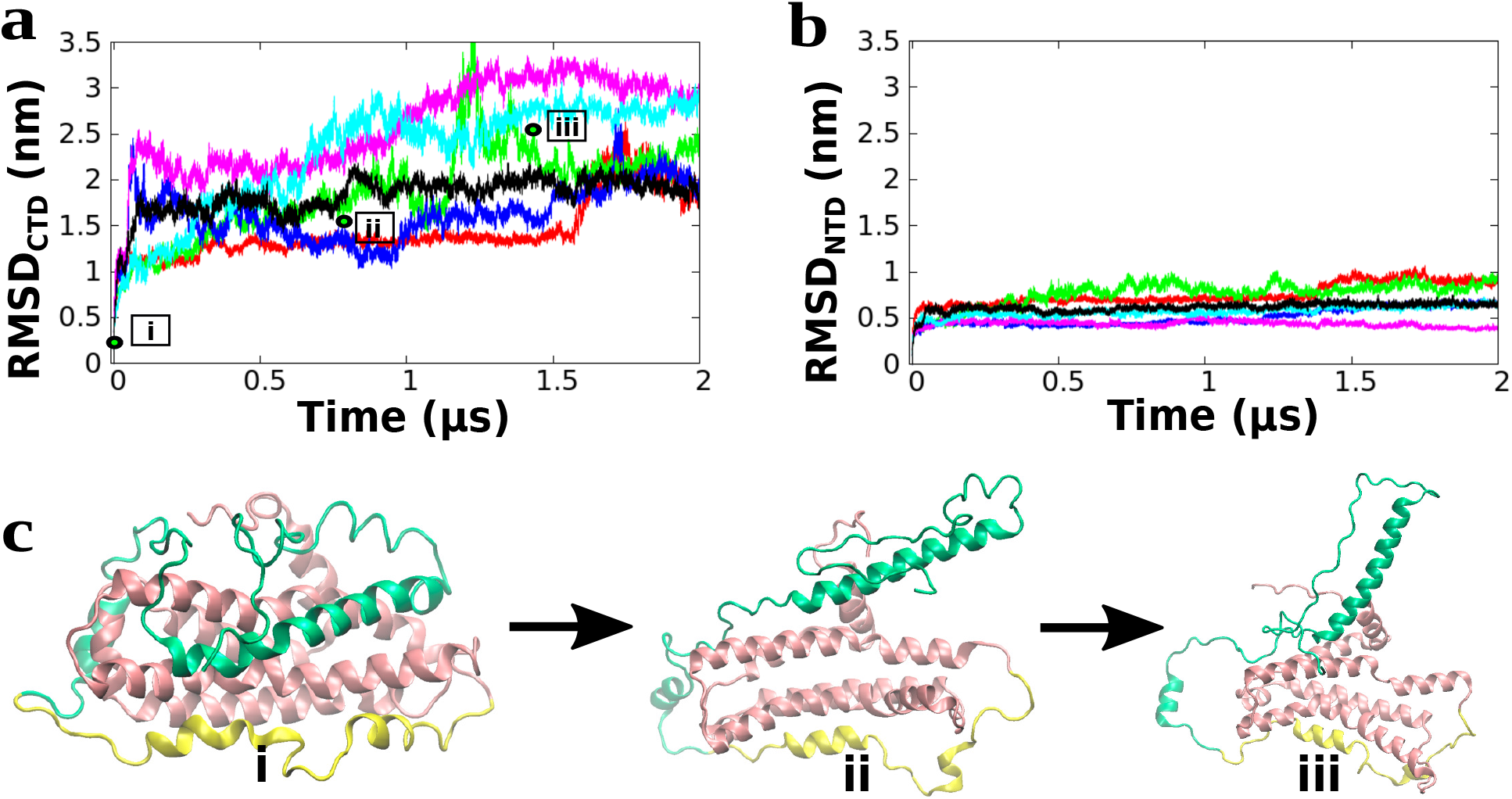
**a.** The time series of RMSD of C-terminal domain (CTD) relative to NMR-derived structure of apoE3 (pdb: 2l7b)^30^ (initial structure of the MD simulation) indicating large opening of the C-terminal domain(CTD). **b**. The time series of RMSD of N-terminal domain (NTD) indicating considerably weaker lipid-induced conformational change of the NTD within the 2 microsecond time-scale. Different curves in figure a-b correspond to individual simulation trajectories. **c**. The snapshots at three time-points highlighted in part a of the figure. These snapshots indicate the closed CTD structure in the beginning of the simulation, the partly opened CTD structure and the fully opened CTD respectively (Refer to color code in figure 1)

### Lipid binding leads to disruption of apoE3 salt-bridges and Intrinsically disordered regions

Analysis of the computer simulated trajectories provides several key insights into the atomistic details of the structural changes due to lipid-induced unfolding of C-terminal domain.

As shown in Figure S5 we find that the assembly of the lipid particles around the protein mostly retains the helicity of apoE3, except at few regions of C-terminal domains of protein. The subtle changes in helicity in few locations of apoE3 is relatively insignificant, especially when compared to its E4 isoform (i.e. apoE4) investigated in previous study.^46^ All in all, the secondary structures of major part of apoE3 remain unaltered even in presence of the lipid.

Inspection of the simulation trajectories reveal that the lipid assembly around the protein disrupts several salt-bridge interactions between N and C-terminal domains. The presence of these salt-bridges had been earlier reported by Chen et al^30^ as one of the stabilising factors of monomeric apoE3. As shown in figure 4a, there are five main salt-bridges establishing contacts between C-terminal domain and helices 2/3 of N-terminal domain in aqueous medium: salt-bridge region labelled 1 is a pair of salt-bridges closely packed together consisting of 217Arg-50Glu and 213Arg-49Glu; similarly other salt bridges are characterised as follows 2:233Lys-132Glu, 3: 238Glu-114Arg and 242Lys-110Asp, 4: 255Glu-95Lys and finally 5: 262Lys-88Glu. Our simulation confirms that these salt-bridges remain mutually intact in absence of lipids (see left panel in figure 4a). However, analysis of our simulation trajectories in presence of lipids predicts that the salt-bridges start loosing their stability in presence of phospholipids. Especially, the tendency of separation of C-terminal domain increases as we spatially move from salt-bridge 1 to 5, highlighting the fact that the C-terminal domain has a higher propensity to loose contact from N-terminus domain at region 5 than at the region 1. Figure S6 establishes this varying tendency of the different salt-bridges to open-up. The Salt-bridge 5 having a peak at larger distance signifies that it has a higher opening tendency and as we move towards salt-bridge 1, this propensity continues to decrease.

**Figure 4:**
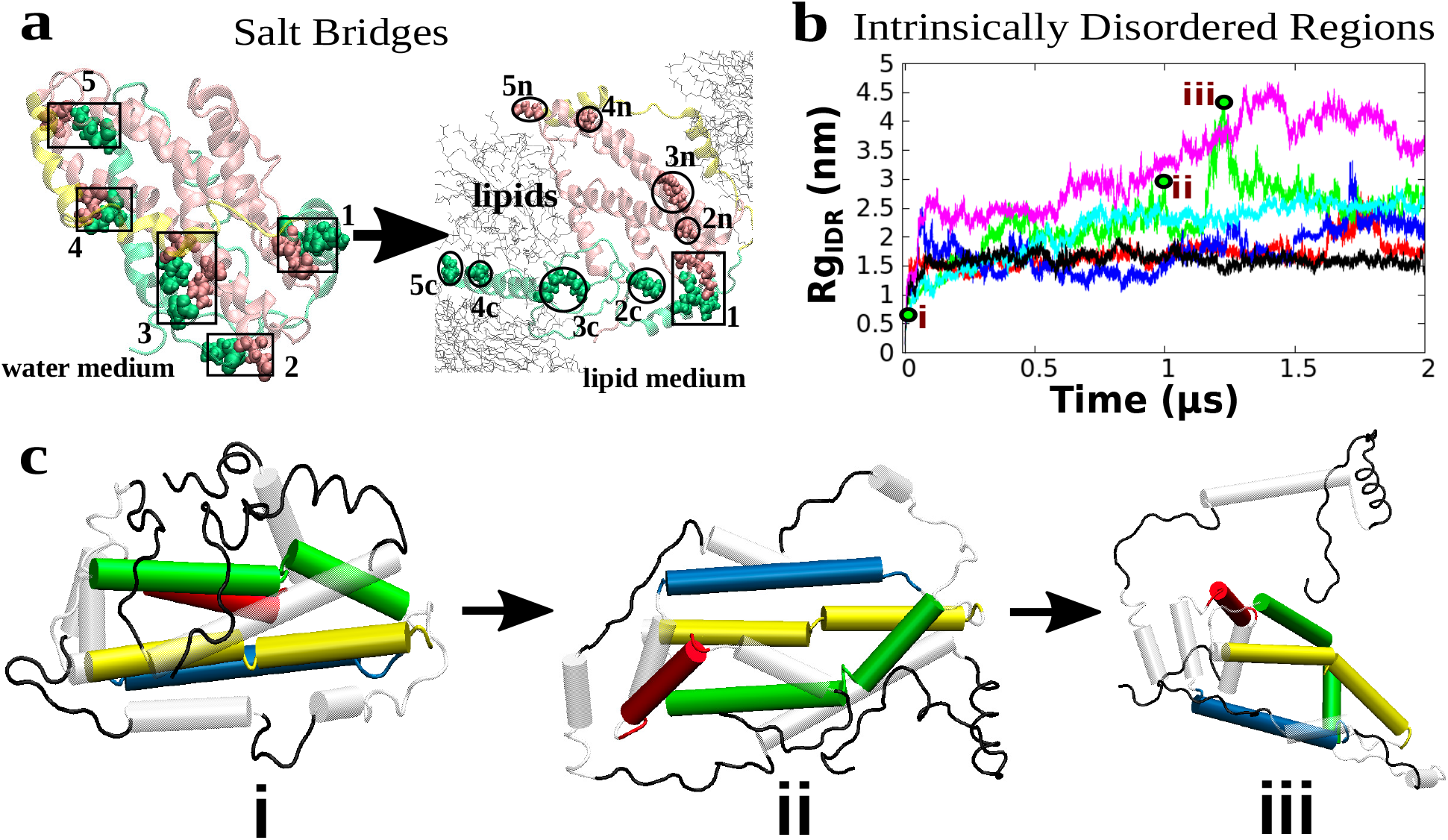
The effect of lipid-assembly on apoE3 structure: **a**. The disruption of salt-bridges 1-5 due to exposure to lipids. In the lipid medium 5n,4n,3n and 2n regions are components of salt bridge from NTD while 5c,4c,3c, and 2c correspond to the CTD components. Saltbridge-1 majorly remains intact. **b**. Time series of radius of gyration of the intrinsically disordered regions (IDR) of the protein for different trajectories. Different curves correspond to individual simulation trajectories. **c**. The snapshots at multiple time-points highlighted in the part b of the figure.

The role of intrinsically disordered regions (IDR) (residues 1-11, 181-187, 198-209, 224-235, 264-299) in stabilising the tertiary fold of apoE3 has been previously emphasised by structure-based investigations.^31^ Our current simulation trajectories involving lipid particles and apoE3 reveal that, upon assembly of lipids around the protein, these IDRs gradually move away from the N-terminal helix bundles. The IDRs of the protein tend to loose contact with the bundle of four helices of the N-terminal domain which is reflected in the time profile of the collective radius of gyration (figure 4b). The progression of increasing radius of gyration of the IDRs (figure 4b) and the accompanying snapshots at different time points (figure 4c) graphically quantify the overall process of dissociation of IDRs upon the unfolding of C-terminal domain.

### Wild type sequence of apoE3 display similar propensity of lipid interaction

To check for robustness of observed mechanism of lipid-induced conformational change in this monomeric mutant sequence of apoE3, we perform additional simulations involving wild-type form of apoE3. Towards this end, we backmutated five residues in NMR structure of monomeric mutant^30^ and computationally designed a wild-type form of the apoE3 (F257/W264/V269/L279/V287). Subsequently we had carried out independent long MD simulations, starting with in-silico designed wild-type apoE3 in water and in similar concentration of aqueous solution of DPPC as employed in the case of the mutant. Figure 5 compares the equilibrated snapshot of the wildtype monemeric apoE3 with that of monomeric mutant in neat water and in aqueous DPPC media in two independently performed simulations . We find that the conformation of the protein in wild-type form resembles very well to that of monomeric mutant in both neat water and lipid-bound state: the wild type protein, otherwise structurally stable in water, opens up in presence of lipid via a very similar mechanism described earlier in case of mutant. This result is consistent with previous experimental demonstration^34,35^ that the structure and stability of the monomeric mutant are identical to wild-type apoE3 and both display similar lipid-binding activities.

**Figure 5:**
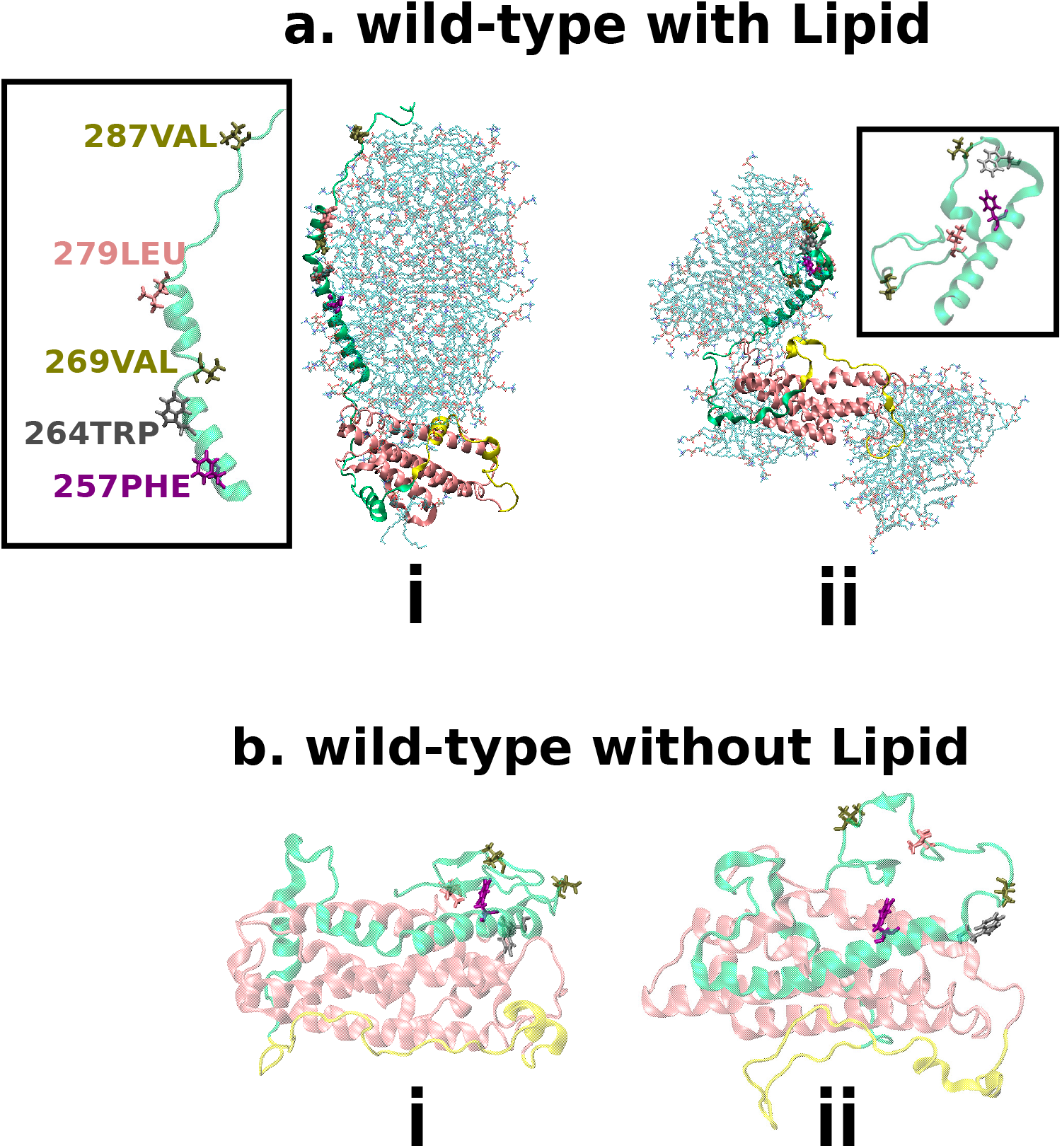
**a.** The representative snapshots of wild type sequence of the apoE3 (F257/W264/V269/L279/V287) obtained in two independent all-atom simulation trajectories with simulation box containing 135 molecules of DPPC along with the water molecules (water not shown). **b**. The structures attained in two independent all-atom trajectories of wild type sequence of the protein in neat water (i.e. without any lipids molecules). Both types of simulations were run for at least 1.5 *µs*. (refer to the color code described in the figure 1).

## Slow lipid-induced unfolding of N-terminal domain as the final stage

### Coarse-grained simulation captures slow unwinding of N-terminal domain

Results in the previous section showed that our multi-microsecond long atomically detailed simulations were able to capture the lipid-induced reorganisation process of C-terminal domain of apoE3. However, N-terminal domain of the proteins remained largely unperturbed by the lipids within the micro-second long time scales of the all-atom simulations. Accordingly, we inferred that any further conformational organisation of the apoE3 might require exploring the conformational phase space for a longer time scale than that covered during the time period of all-atom simulation. Modelling with coarse-grained force-fields allows one to reduce the number of degrees of freedom and choosing a larger time-step to sample the conformational landscape more extensively. This prompted us to adopt a coarse-grained model of the protein-lipid-water mixtures so as to facilitate the access of relatively longer simulation time scale. Towards this end, the equilibrated configuration as obtained from the all-atom MD simulations at the end of 2 *µs* long simulation, in which C-terminal domain has been completely separated from the N-terminal domain (see snapshot in figure 2a), served as the initial conformation for the subsequent coarse-grained simulation. We employed MARTINI force field^43–45^ to map the atomistic conformations of apoE3, DPPC lipids and water into their respective coarse-grained representations. Our all-atom investigation suggested that the secondary structures of apoE3 is mostly retained in neat water as well as in presence of lipid particles, which justified a seamless mapping of all-atom lipid-induced conformation protein to Martini coarse-grained model (which otherwise does not allow for change of secondary structure). Figure 6a represents the initial conformation of protein-lipid complex (with C-terminal domain separated) mapped in coarse-grained representation. Supplemental movie S2 and movie S3 depict the dynamical process of conformational reorganisation, as observed in two individual 55 *µs*-long independent coarse-grained trajectories. We find that all helices of N-terminal domain remained fairly closed to each other for a long part (around 20-25 *µ*s) of coarse-grained simulation time, while the C-terminal domain remained separated. However, after sampling the ensemble of close-packed N-terminal domain conformation for a long period of time, intriguingly helix1/helix2 of N-terminal domain started getting separated from helix3/helix4 and the lipid molecules start invading the resulting intermediate gaps between these two sets of helices of N-terminal domain.

**Figure 6:**
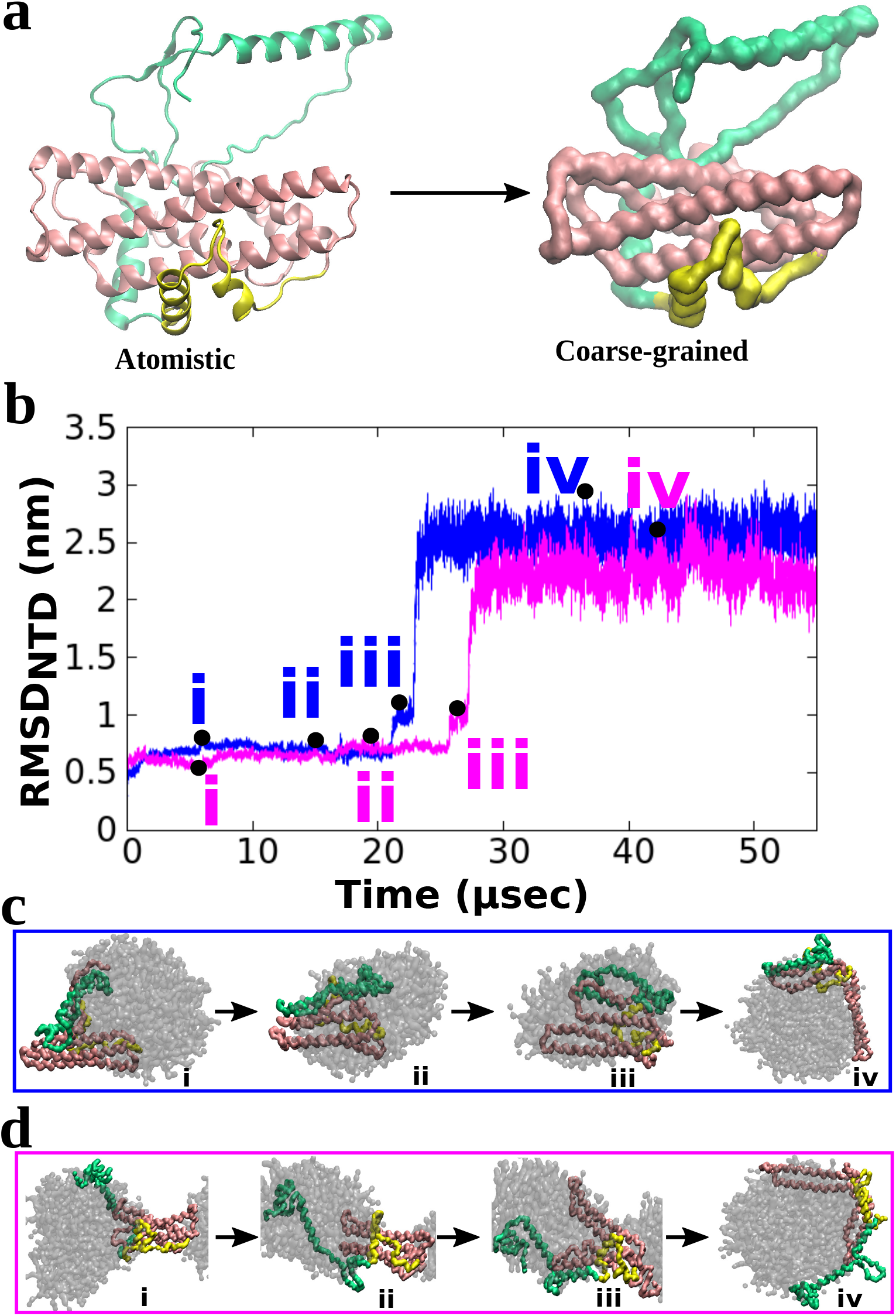
**a.** Coarse-graining of all-atom conformation of apoE3 (obtained at the end of 2 *µs* long apoE3-lipid all-atom simulation) for exploring long time scale apoE-lipid interaction behaviour. **b**. The time series of RMSD of N-terminal domain (NTD) with respect to the initial conformation of the coarse-grained MD simulation. Different curves correspond to individual simulation trajectories. **c**. Four important snapshots from trajectory-1. **d**. Four important snapshots from trajectory-2. In both these trajectories, the NTD open up with the helices 1-2 separate apart from helices 3-4. (refer to the color code described in the figure 1)

To characterise the lipid-induced conformational organisation of the N-terminal domain, we compute the time profile of RMSD of N-terminal domain during the period of coarse-grained simulations (figure 6b). We find that in both trajectories, each spanning 55 *µs*, the RMSD of N-terminal domain first remained low (figure 6b) for a long duration of the simulation until 20-25 *µs*. Subsequently, beyond 20-25 *µs* long simulation period, we find that the RMSD of N-terminal domain shows a sharp increase in its value, thereby suggesting a sudden conformational organisation of N-terminal domain. After the increase at 20-25 *µs*, the RMSD profiles in figure 5b plateaued beyond 30 *µs* and remained unchanged for the remainder of 55 *µs* trajectories. Parts c and d of figure 6 represent the key conformations from various time point of these two coarse-grained simulation trajectories: the gradual lipid-induced opening of N-terminal domain is quite evident in these snapshots from these trajectories. Both profiles indicate that, starting with conformation similar to snapshot I (see figure 6), helix1/helix2 and helix3/helix4 and N-terminus domain unwraps in stage-wise manner. Snapshot II and Snapshot III shown in figure 6 for both trajectories are characterised as the early-stage and late-stage intermediate conformations seeding the unwrapping process of the N-terminal domain in both the trajectories. Finally, snapshots IV in the bottom panels of the figure 6 c-d represent the final equilibrated conformation of the lipoprotein particle, as obtained in the current investigation.

In a bid to dissect the key residues mediating the unravelling process of N-terminal domain, we probe the distance between the centre of mass of residues 45-54 and residues 124-129 (hereby called *D*1), with the progression of simulation. These two residue-sets (coloured black in figure 7) act as the linker between N-terminal helices 1/2 and helices 3/4 respectively. As shown in figure 7, *D*1 i.e. the distance between the linkers mostly remained confined within 1 nm until 20-25 *µs*, indicating reversible exploration of closed conformation by the ensemble of apoE3-lipid complex. However, in both the trajectories the distance between these two linkers show a large jump to a value of 8.5 nm, signalling the structural reorganisation of the N-terminal domain. On the same figure, the minimum distance between C-terminal domain and helix 3/helix 4 of N-terminal domain (referred as *D*2) is also plotted. We find that the C-terminal domain (shown in green) initially rests on the helix 3/helix 4 of the N-terminal domain (shown in pink). However, with increased residence of lipid molecules in contact with the helices, the distance between C-terminal domain (especially set of non-polar residues in 227-240 and 280-298) and helices 3/4 starts gradually increasing (figure 7, right axis). The overlay between these two distances i.e. *D*_1_ and *D*_2_ in figure 7 clearly indicates that the separation process of helices 1/2 from helices 3/4 is strongly correlated with the separation of C terminal domain from helix3/helix 4 of N-terminus domain. The helices 3 and 4 have been marked in each of the two final snapshots.

**Figure 7:**
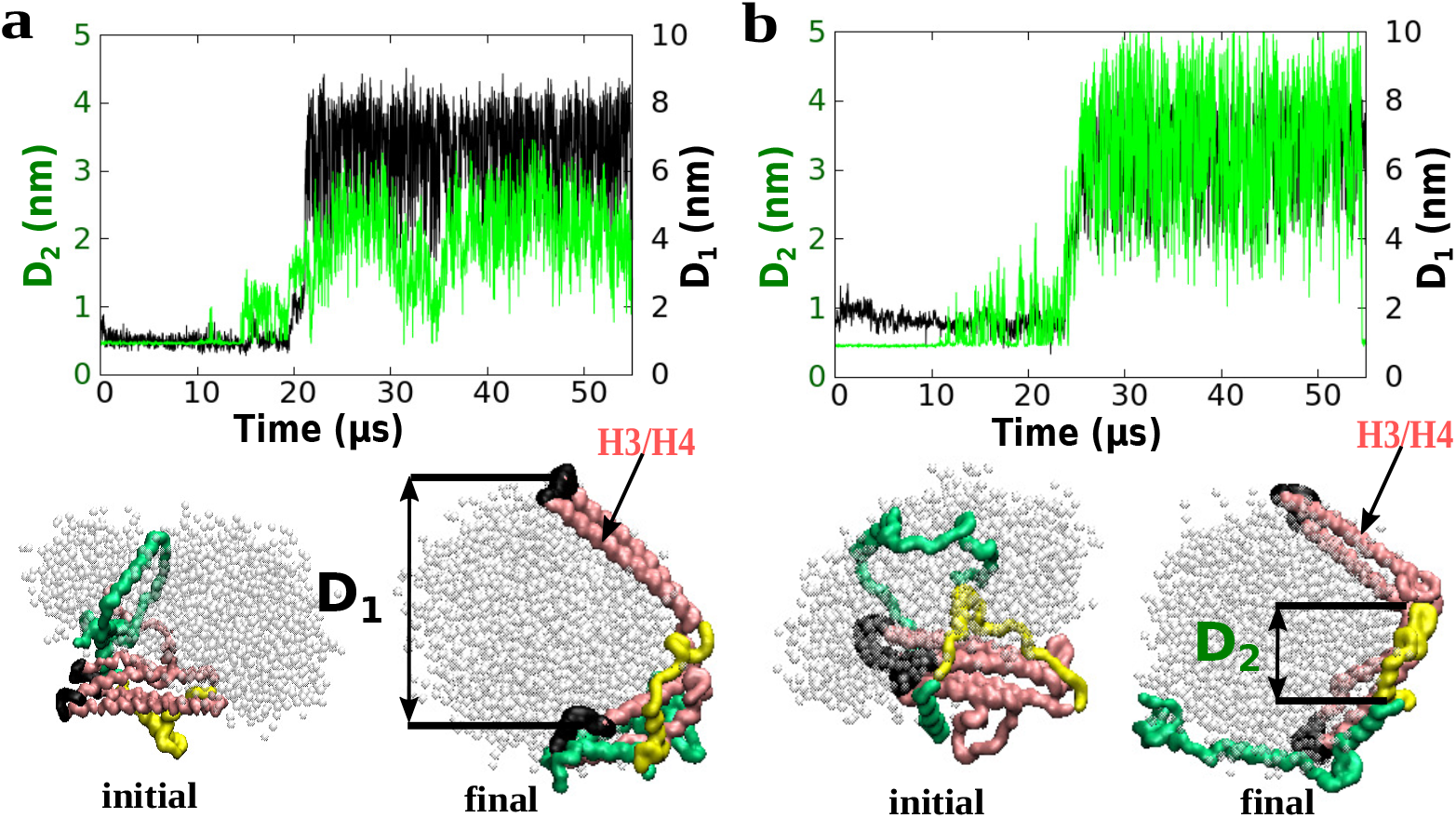
Dissecting the key determinants of lipid-induced unwinding of N-terminal domain in coarse-grained simulation: **a**. For trajectory 1: Right hand axis: The time series of the distance (black curve, referred as *D*1) between the two linker regions (shown in black in snapshot below) of the N-terminal domain (one between the helices 1/2 and the other between helices 3/4) . Left hand axis: The time series of the minimum distance between the C-terminal domain and helices 3/4 shown (green curve,referred as *D*2). **b**. same as in part a but for trajectory 2. Bottom: Shown in the bottom the initial and final structure in each of the trajectories. The extent of opening up of the N-terminal domain has been pointed out by the double-ended-arrow line in the final snapshot for each of the two trajectories.

### Comparison of simulated apoE3.lipid complex morphology with experiments

The final open structure, represented in figure 7 is majorly stabilised by the strong hydrophobic association between lipids and reorganised N-and C-terminal domain. The ultimate question is how realistic is the morphology of the lipoprotein particle, as obtained from our simulation? While changes in secondary structure can not be modelled within the coarse-grained model, changes in tertiary structures are unrestricted and in principle realistic within the general approximations (See reference^47^ and references of examples there-in). Nonetheless, to check the sanity of the apoE3-lipid complex, as obtained from the coarse-grained simulation, we undertook an existing strategy proposed by Wassenaar et al^48^ to reverse-map the structure to its atomistic resolution. The reverse-mapped all-atom structure, thereby obtained, were subsequently subjected to equilibration via MD simulations. Figure 8 depicts the final snapshot of the all-atom representation of the structure. Even though the structure obtained from the current investigation were obtained from a coarse-grained model, It was gratifying to find that an inverse-mapped all-atom representation, retains similar conformation, thereby lending credence to the proposed ‘open conformation’.

**Figure 8:**
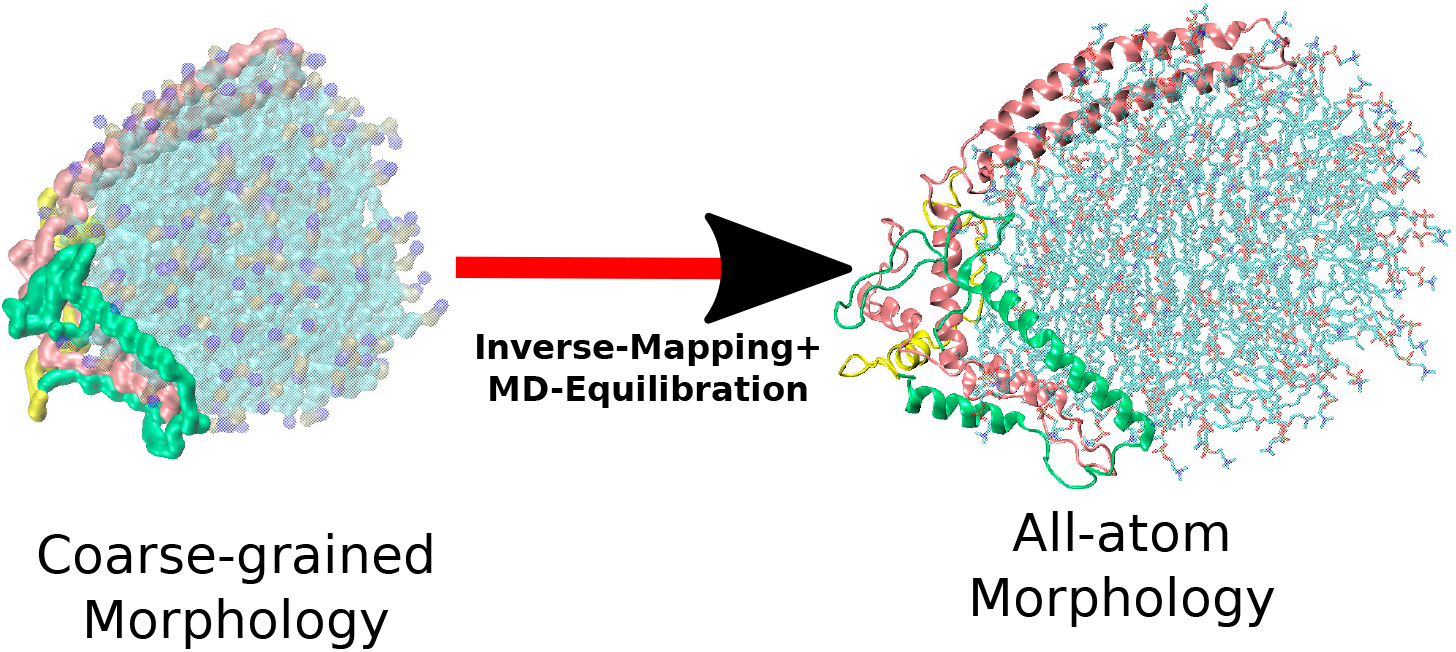
All-atom realisation of the apoE3-lipid complex obtained after reverse-mapping (and subsequent equilibration) of the complex which was derived from coarse-grained simulation.

There are large volume of precedent experimental investigations based on crystallography and SAXS which have proposed a diverse range of morphologies for apoE3-lipid complex.^4,22,23,49^ The morphology of the resultant complex, as obtained from the current work shows very close resemblance with an earlier proposal of a ‘open structure’.^4^ There is also report of a similar open model^12^ for apoE3-lipid complex, where a structure of apoE3 N-terminal Domain, as having the N-terminal helical bundle opened up without any observable helical structure disruption has been proposed. In this regard, Weisgraber and co-worker^11^ provided a detailed investigation of the surface properties of N-terminal domain where a similar open structure is reached with four-helices bundle in N-terminal domain reorganised as the pink portion in the figure 9c. The final structure attained at the end of current coarse-grained simulations conforms to these proposed models from experiments. To the best of our knowledge, the current work is the first to provide a higher resolution molecular picture of the apoE3 lipoprotein particle and to capture the complete molecular mechanism of the complexation process of full protein in structurally precise and time-resolved manner.

**Figure 9:**
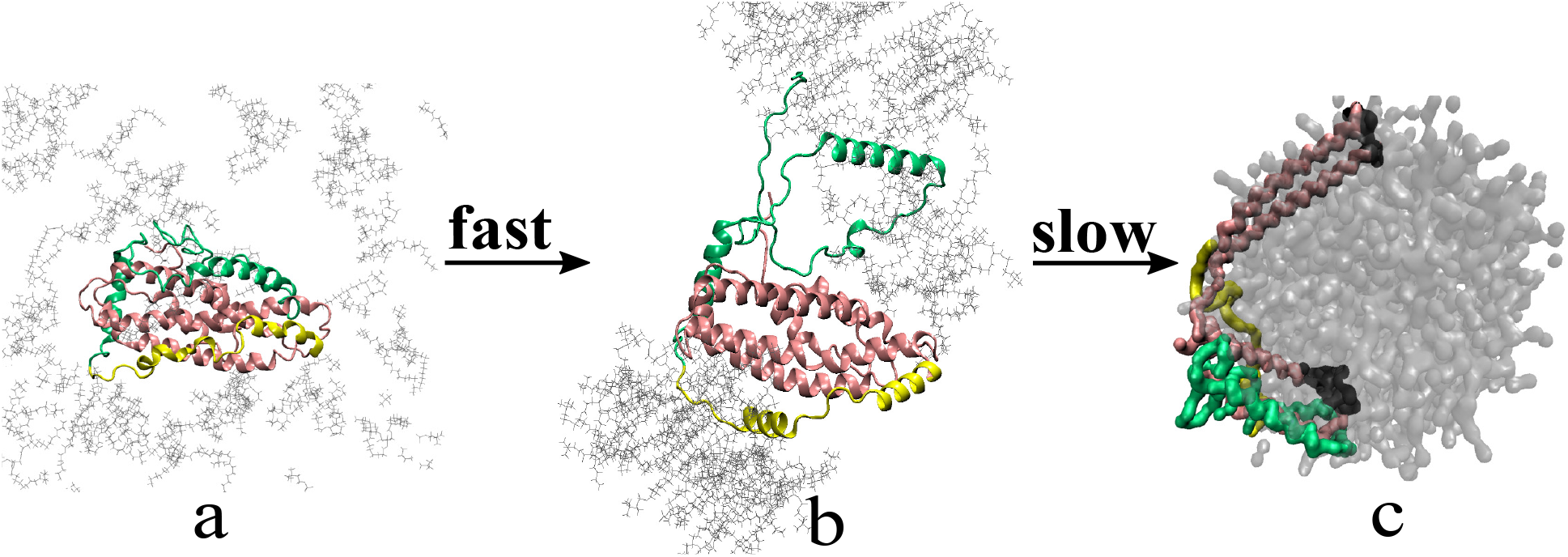
The representative snapshots summarising the overall two-stage opening of apoE3 in presence of phospholipid environment: The atomistic simulations captures the opening up of the green colour C-terminal domain which is a fast process. The slow opening up of the N-terminal domain shown in pink colour is captured in coarse-grained simulation.

## Discussion

The full complexation process of apoE3 and phospholipids, as elucidated in current work’s unbiased MD simulation efforts is summarised in figure 9. The simulations reported in the current work elucidate a two-stage reorganisation process of apoE3 tertiary fold in presence of lipids : i) a fast lipid-induced opening of C-terminal domain of the apoE3 from the rest of the protein (as observed in our all-atom simulations), followed by ii) a slow lipid-mediated inter-helix separation within N-terminal domain (as observed in our long coarse-grained simulations). The present work’s detailed molecular-level kinetic illustration of two-stage conformational reorganisation of apoE3 in presence of phospholipids, provides solid ground to earlier proposals arising from biochemical experiments ^25,28,29^ and structure-based hypothesis^30,31^ that the complete lipid binding process is unlikely to occur in a single stage.

The observation of preferential binding of lipids to the C-terminal domain of apoE3 in the current simulation is consistent with proposed structural basis of lipid binding in apoE3 by Chen et al^30^ and kinetic studies by Garai et al.^26,50^ In particular, the simulation identifies residues 243-296 as the prominent lipid binding locations in C-terminal domains. Frieden and coworkers have previously hypothesised about the important role that IDRs and salt-bridges potentially play in maintaining the tertiary fold between N-terminal domain and C-terminal domain of apoE3 in water.^31^ The current work quantitatively demonstrates that the lipid-induced disruption of several salt-bridges between N-and C-terminal domains and outward movement of IDRs drive the separation of C-terminal domain from the rest of apoE3. The uncoupling process of helices in N-terminal domain, as observed in our coarse-grained simulations, is found to be slow and mainly guided by weakening of interactions between helix 3/helix4 of N-terminal domain and hydrophobic residues of C-terminal domain by the intercepting lipid molecules. Additionally, the simulations demonstrate that the lipid-induced conformational organization in mutated apoE3 is quite robust in wild-type as well. The final open structure of the lipoprotein particle, represented in figure 9c is majorly stabilised by the strong hydrophobic association between lipids and reorganised N-and C-terminal domain. There are large volume of precedent experimental investigations based on crystallography and small-angle X-ray scattering techniques which have proposed a diverse range of morphologies for apoE3.lipid complex.^4,22,23^ However, many of the previous models had been proposed by drawing parallel with the previously proposed model for more popular apoAI-lipid complexes.^25,51^ Many of the models propose dimer of apoE3 to assemble together with lipids. However, more recent conclusion by Garai et al^26^ that apoE3 binds to lipid in monomeric form indicates that any proposal of the apoe3.lipid complex needs to be based on the monomeric apoE3. Towards this end, the current investigation, which is based upon the same, provides a physical ground to the possible morphology of apoE3.lipid complex. The resemblance of morphology of the complex as obtained from the current work with an earlier proposal of a ‘open conformation’ ^4^ provides a solid ground to the step-wise unfolding of C-terminal and N-terminal domain of the protein.

What might be the functional implication of lipid-induced conformational change of apoE3, as derived in the current investigations? Prior hypothesis suggests that only in the maximal lipid-associated state does apoE3 become adept to act as a ligand for LDLR,^11,16,23,30^ following which the lipid-transport metabolic circuit becomes effective. The two-step lipid complexation mechanism of apoE3, namely fast C-terminal domain unwinding and subsequent slow N-terminal inter-helix separation, as elucidated in the current work, rules out any possibility of partially lipid-bound or lipid-free apoE3 in its native tertiary fold, to be acting as ligand for LDLR. This pre-requisite of optimal lipid-induced conformational re-organization of apoE3 leading to an ‘open conformation’ for LDLR recognition, has also been proposed in prior experimental investigations including FTIR spectroscopic investigations,^12^ fluorescence resonance energy transfer-based experiments,^52^ tryptophan fluorescence quenching studies.^11,16^ Overall, present data obtained from MD simulation support the view that a lipid binding-induced conformational adaptation of apoE3, is an essential feature of apoE function as a ligand for receptor-mediated endocytosis of plasma lipoproteins.

## Conclusion

In conclusion, we provide below a summary of the salient observations made and insights drawn from this investigation:

- The simulation unequivocally shows that conformational state of apoE3 individually in water and in lipid environments are mutually distinct: while in water the protein remains conformationally globular, it undergoes significant conformational reorganization process in presence of lipid molecules. Together, the simulation characterizes the lipid-induced reorganized state of this key protein.
- The investigation demonstrates that it is the predominantly large segment of the C-terminal domain of the apoE3, where lipid molecules make their first localized contact with the protein. The simulation also statistically identifies the key residue segments in C-terminal domains where the lipid molecules first settle in.
- The simulations reported in the current work in particular elucidate a two-stage reorganisation process of apoE3 tertiary fold in presence of lipids : a) a fast lipid-induced opening of C-terminal domain of the apoE3 from the rest of the protein (as observed in our all-atom simulations), followed by b) a slow lipid-mediated inter-helix separation within N-terminal domain (as observed in our long coarse-grained simulations).
- The current work quantitatively demonstrates that the lipid-induced disruption of several salt-bridges between N-and C-terminal domains and outward movement of IDRs drive the separation of C-terminal domain from the rest of apoE3. The uncoupling process of helices in N-terminal domain, as observed in our coarse-grained simulations, is mainly guided by weakening of interactions between helix 3/helix4 of N-terminal domain and hydrophobic residues of C-terminal domain by the intercepting lipid molecules.
- We show that the lipid-induced conformational activated state remains robust both in an experimentally established monomeric mutant and the wild-type version of this protein.
- Finally the eventual apoE3-lipid complex, as obtained from coarse-grained simulation, was found to retain the morphology in an refined all-atom representation of the complex.

## Author Contribution

JM and DP designed the project. DP performed the computer simulations. JM, DP analysed the results and wrote the paper.

## Supplemental Information

Supporting information with figures comparing snapshots of apoE3 in presence and absence of lipids in multiple trajectories, the time-series of number of C-terminal domain crowding around the N-terminal domain, comparison of lipid-induced morphology across multiple version of Gromacs, comparison of temporal evolution of the lipid assembly by turning of or turning on the dispersion correction term, residue-wise helicity of apoE3 in presence and absence of lipid molecules, quantification of lipid-induced salt-bridge opening and description of movies.

## Acknowledgements

JM thanks Dr. Kanchan Garai for useful discussions on precedent investigations on the ApoE-lipid interaction. This work was supported by computing resources obtained from shared facility of TIFR Centre for Interdisciplinary Sciences, India. We acknowledge support of the Department of Atomic Energy, Government of India, under Project Identification No. RTI 4007. JM acknowledges Ramanujan Fellowship and Core Research grants provided by the Department of Science and Technology (DST) of India (CRG/2019/001219).

