## Supplementary figures for "Conformational Reorganisation of Apolipoprotein E Triggered by Phospholipid Assembly"

### Supporting Information for "Conformational Reorganisation of Apolipoprotein E Triggered by Phospholipid Assembly"


**Description of Movie:**

**Movie S1:** This Movie depicts the first stage of reorganisation of apoE3 tertiary fold in presence of DPPC lipids (water not shown for clarity). The movie is a representation of an all-atom molecular dynamics simulation trajectory in which DPPC lipids (black) assemble around apoE3 and induces the movement of C-terminal domain (green) away from the rest of apoE3 fold (pink N-terminal domain and yellow hinge region).

**Movie S2:** The movie depicts the second and final stage of phospholipid-induced reorganisation of apoE3 tertiary fold. The movie is a representation of 30 microsecond-long coarse-grained molecular dynamics simulation trajectory in which lipid molecules (black) mediate the slow unwinding of helices 1/2 of N-terminal domain from helices 3/4 of N-terminal domain (pink)

**Movie S3:** A separate coarse-grained trajectory demonstrating the same end-result as in movie S2. The color code is same in movie S2

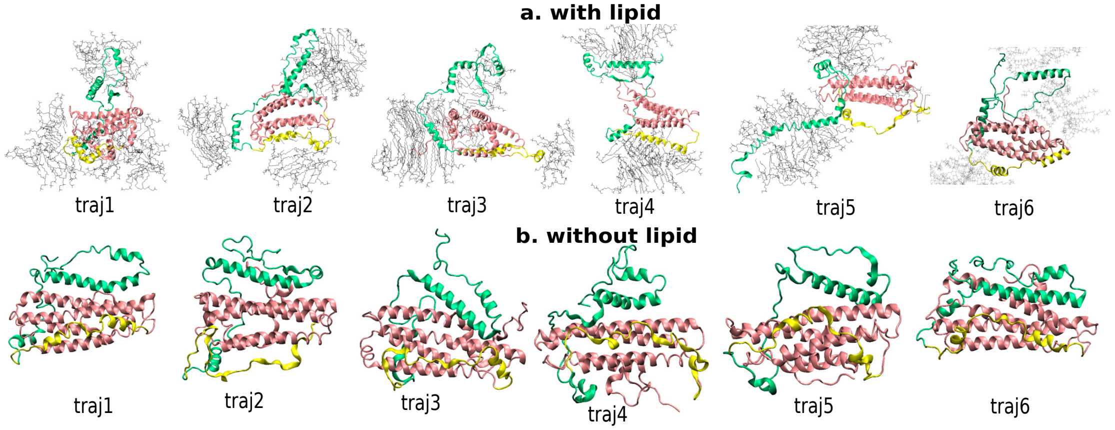


Figure S1: **a.** The representative snapshots obtained in six independent all-atom trajectories with simulation box containing 135 molecules of DPPC along with the water molecules (water not shown). **b.** The structures attained in six independent all-atom trajectories without any lipids molecules. Both types of simulations were run for similar length. (refer to the color code described in the figure 1)

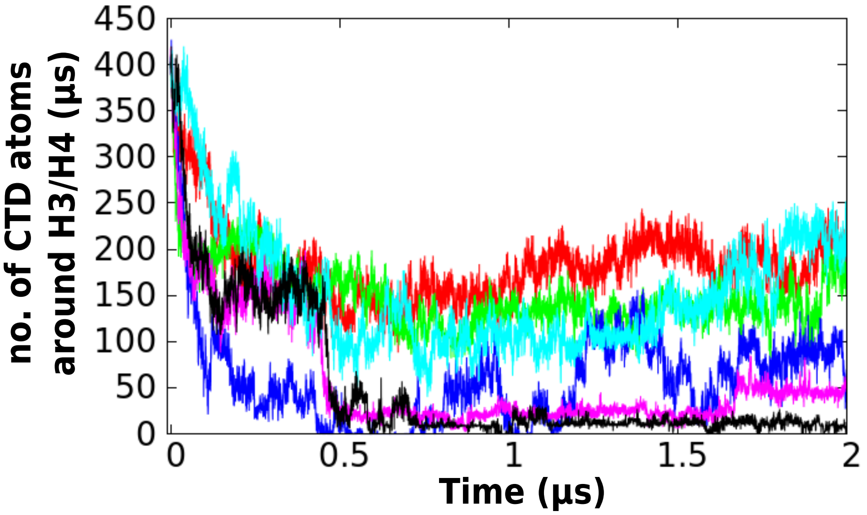


Figure S2: The time series of the number of atoms of C-terminal domain (CTD) crowding around the helices 3 and 4 of N-terminal domain as obtained from all-atom simulation

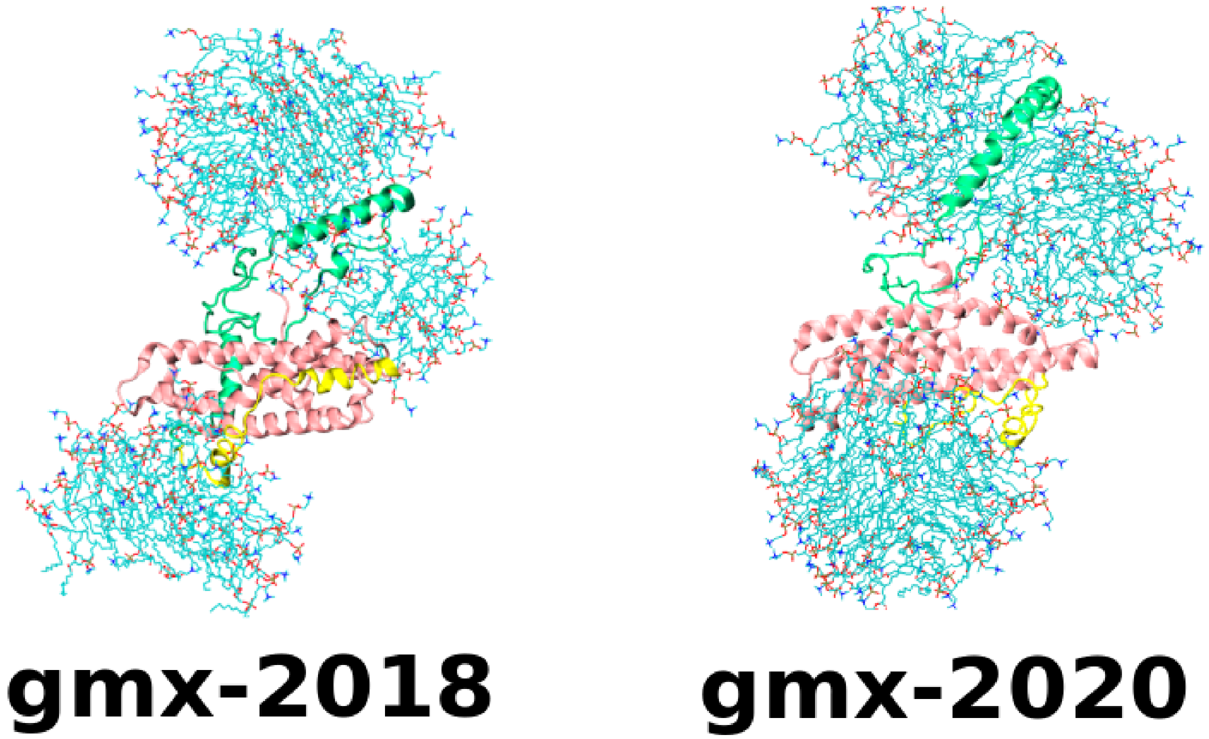


Figure S3: The lipid-assembled morphology of the apoE3 in which C-terminal domain is separated is robust across different Gromacs version (2018 vs 2020) in all atom simulation.

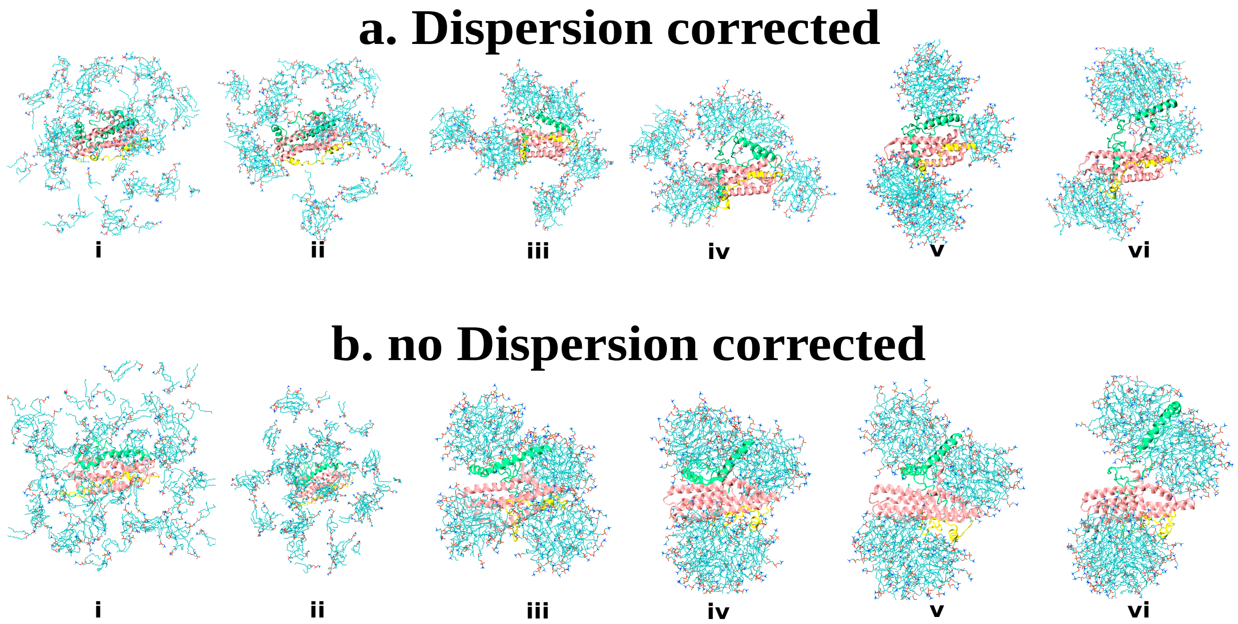


Figure S4: Comparison of evolution of protein/lipid assembly process in presence of dispersion correction, with that in absence of dispersion correction while modelling nonbonding interaction

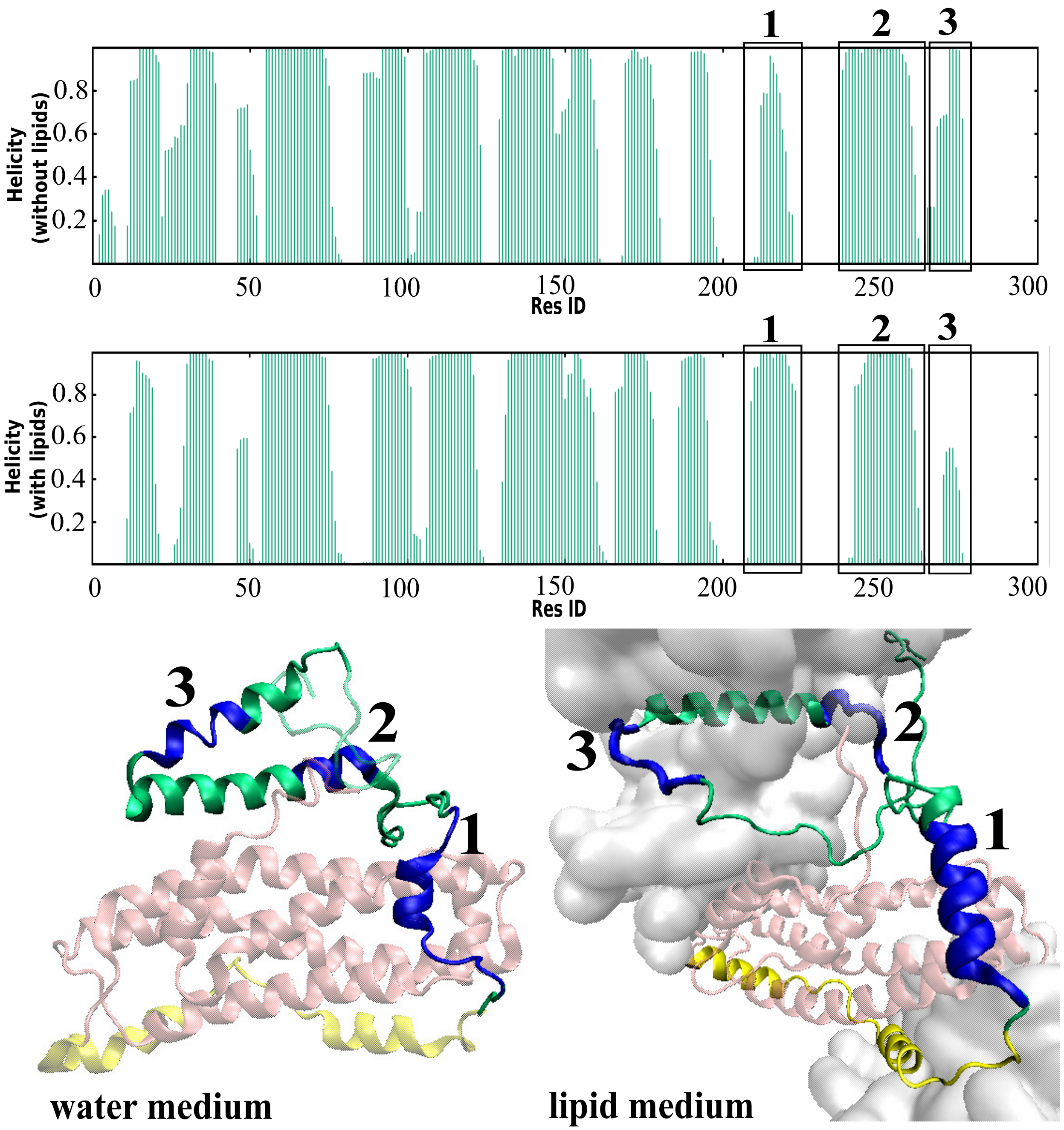


Figure S5: **a.** The helicity of the regions 1, 2 and 3 (marked by boxes on the distribution plot) of the protein is seen to be changing when the aqueous environment of the protein is changed to lipid environment.

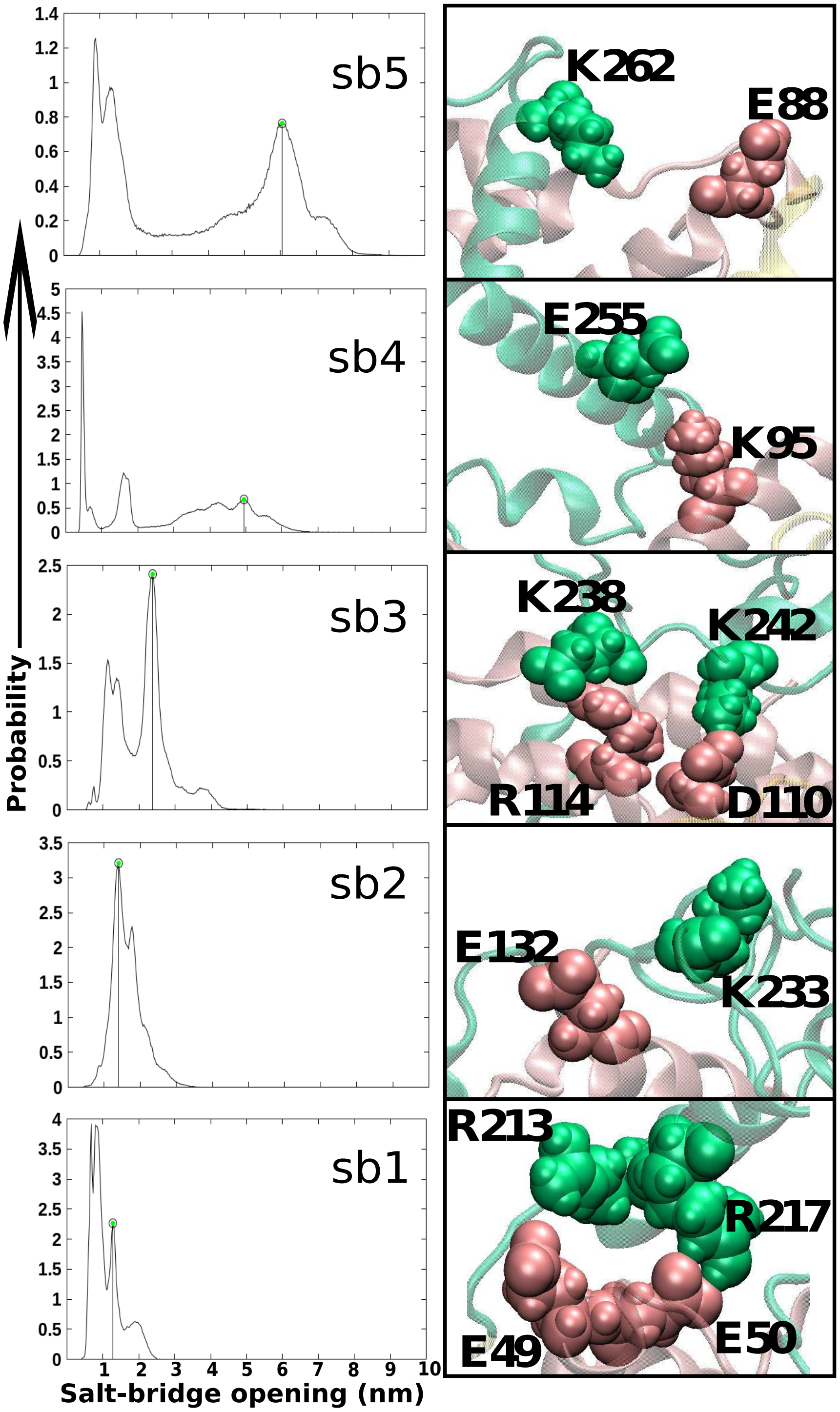


Figure S6: In the left, we have a plot where the x-axis has the salt-bridge opening which is measured by the distance between the centres of mass of the green bubble regions and pink bubble regions in the adjacent snapshot and the y-axis denotes the percentage probability of the occurrence in a trajectory. There are 5 such plots, one for each of the 5 salt-bridges. The salt-bridge 5 has an additional peak at a large distance value of around 6 nm, salt-bridge 4 around 5 nm and so on. Thus we see that salt-bridge 5 has a high tendency to remain open while this tendency gradually reduces as we move towards salt-bridge 1 from salt-bridge 5. In the right panel we have the close-up snapshot of the corresponding salt-bridge.
